# Amoebicidal Action of Licorice-Derived Compounds: Mechanisms and Effects Against *Acanthamoeba castellanii*

**DOI:** 10.1101/2025.02.17.638758

**Authors:** Lijun Chen, Dai Dong, Wenwen Jing, Qingtong Zhou, Meng Feng, Xunjia Cheng

**Author notes:** **Corresponding author**: Author name: Xunjia Cheng, Mailing address Author name: Meng Feng, Mailing address. These authors contributed equally to this work (Lijun Chen, Dai Dong).

## Abstract

**Background:** *Acanthamoeba* is a widely distributed unicellular eukaryotic organism, causing *Acanthamoeba* keratitis, granulomatous amoebic encephalitis and chronic infectious ulcers. At present, there are no effective drugs available for treating *Acanthamoeba* infections, highlighting the urgent need for the development of novel anti-*Acanthamoeba* therapies.

**Purpose:** This study aims to investigate the inhibitory effect of licorice on *A. castellanii* trophozoites growth and elucidate its molecular mechanisms.

**Methods:** The effect of licorice on trophozoite viability was assessed employing the CellTiter-Glo assay, while the viability of cells was evaluated via CCK-8 assay. Apoptotic cells were identified through Hoechst 33,342/PI double staining and caspase-3 detection. ROS level was measured via flow cytometry utilising DCFH-DA. Mitochondrial dysfunction was analysed with JC-1 and mtSOX Deep Red staining. RNA sequencing and RT-qPCR analysis were conducted to investigate the potential anti-*Acanthamoeba* mechanism.

**Results:** Our results showed that isoliquiritigenin (ISL) and glabridin (GLA) effectively inhibited *A. castellanii*trophozoite growth *in vitro* dose-and time-dependently. ISL and GLA induced several apoptosis features, including Hoechst/PI-positive staining and increased caspase-3 expression. ISL and GLA increased intracellular ROS production, decreased SOD expression and mitochondrial membrane potential and enhanced mitochondrial ROS generation. ISL and GLA effectively prevent host cells from *A. castellanii* trophozoites invasion. RNA sequencing indicated that ISL may modulate the NAD^+^ metabolic process, while GLA may influence the sterol metabolic process. ISL reduced the NAD^+^/NADH ratio, while GLA lowered the levels of 7-dehydrocholesterol in *A. castellanii* trophozoites.

**Conclusion:** These findings suggest that ISL and GLA may serve as the active components of licorice for treating *A. castellanii* trophozoites. Our findings suggest that ISL suppresses trophozoites by regulating NAD^+^ metabolism, while GLA inhibits trophozoites through the regulation of sterol metabolism.

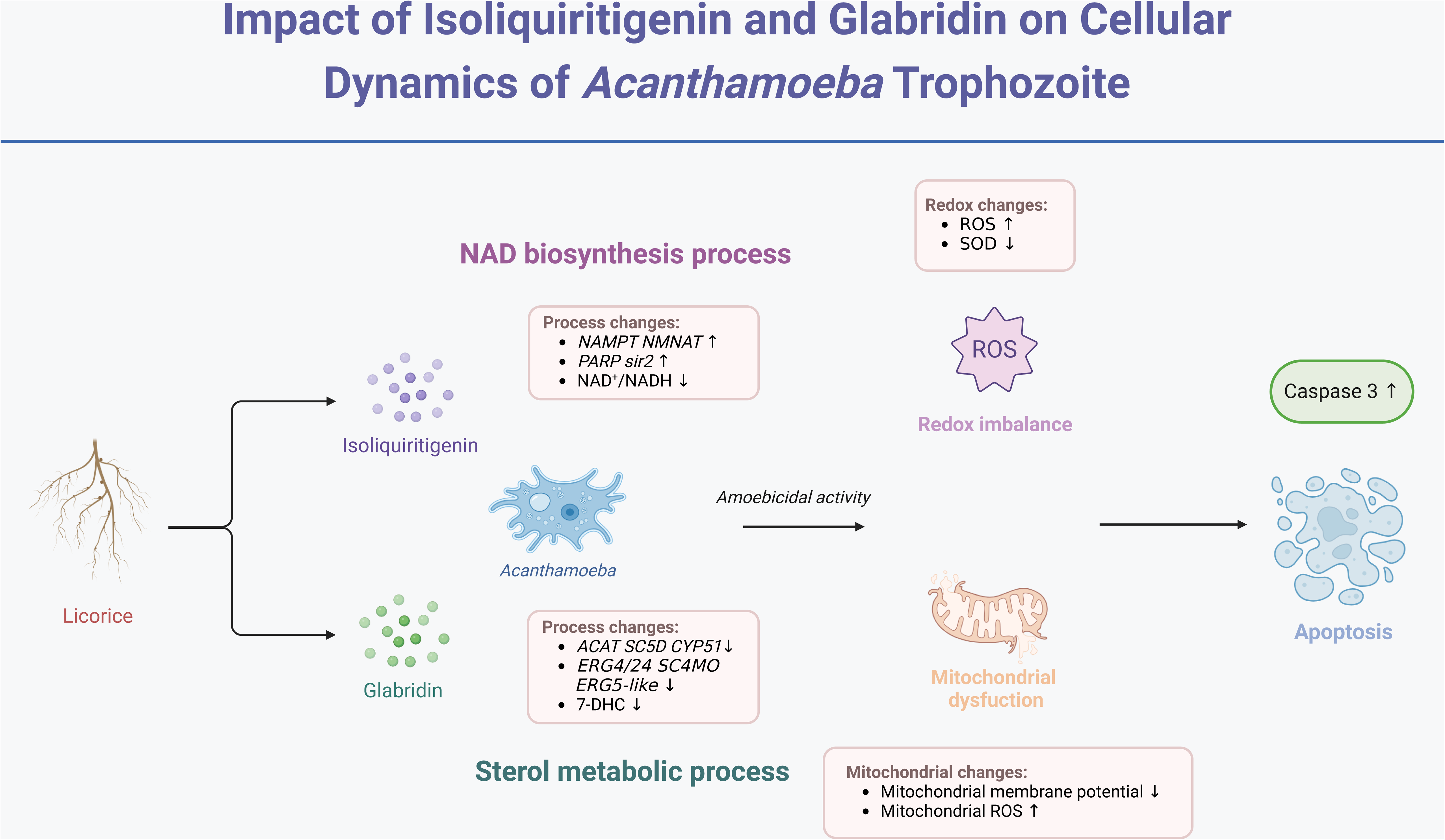

## Introduction

*Acanthamoeba* spp. are free-living amoebae that occur worldwide and are often overlooked as potential sources of infection leading to severe clinical complications in humans. As a significant human pathogen, *Acanthamoeba castellanii* causes various infections, from *Acanthamoeba* keratitis (AK) to granulomatous amoebic encephalitis (GAE) (Rayamajhee et al., 2022)—a rare but a fatal disease. Infectious keratitis typically leads to sight-threatening complications (Aiello et al., 2024). The WHO has classified AK as a neglected tropical disease, emphasizing its significance as a global public health concern. However, the development of effective drugs against *Acanthamoeba* infection remains a critical challenge.

Natural herbaceous plants have been widely used in traditional Chinese medicine for centuries owing to their diverse pharmacological activities, including anti-inflammatory, antiviral, and antibacterial properties. However, systematic research on the specific bioactive compounds and their mechanisms of action remains lacking. Among these compounds, Isoliquiritigenin (ISL) and Glabridin (GLA) are flavonoid-based compounds derived from Licorice. Licorice contains two essential classes of bioactive compounds, triterpenoids and flavonoids, which contribute to its medicinal properties (Wu et al., 2024). ISL and GLA, which are natural flavonoids with a chalcone structure, are extracted from Gancao and exhibit unique and specific biological properties (Gaur et al., 2014; Chen et al., 2018; El-Ashmawy et al., 2018; Wang et al., 2021; Zhang et al., 2022, Yu, 2023; Sun et al., 2023). ISL functions as a dual PPARγ and Nrf2 agonist, exhibiting antiviral and anti-inflammatory properties that reduce lung viral titres and morbidity in mice infected with the PR8/H1N1 virus (Traboulsi et al., 2015). Additionally, ISL effectively suppresses the replication of RNA or DNA viruses *in vitro* (Sekine-Osajima et al., 2009; Wang et al., 2023).

Similarly, increasing evidence suggests that GLA enhances the effectiveness of first-line antibiotics, including norfloxacin and vancomycin, against *Staphylococcus aureus*, suggesting its potential as an adjunct therapy for combating antibiotic resistance (Yang et al., 2024). GLA up-regulates ROS, NO, and MDA levels while reducing SOD expression levels, contributing to its inhibitory effects on *Staphylococcus aureus* (Singh et al., 2015). Furthermore, GLA inhibits the production of inflammatory mediators in HL-60 cells and dendritic cell maturation dose-dependently by blocking NF-κB and MAPK signalling pathways (Kim et al., 2010; Chandrasekaran et al., 2011). Although ISL and GLA have been extensively studied in the fields of inflammation and cancer, research on their anti-parasitic activity remains limited. ISL exhibits a multi-stage anti-malarial effect with synergistic activity against multidrug-resistant *Plasmodium falciparum* (Kumar, 2024). Studies on results and validation in treating *Acanthamoeba* infection remain lacking.

Concurrently, their effects on *A. castellanii* trophozoite growth *in vivo* and the underlying molecular mechanisms remain unclear. For the treatment of intractable free-living amoeba, a comprehensive and multidisciplinary approach should be applied. Thus, this study, as the first, aims to evaluate the effect of ISL and GLA on *A. castellanii* trophozoites and clarify its potential underlying molecular mechanisms.

## Materials and methods

### Amoeba and host cell cultivation

*Acanthamoeba castellanii* (ATCC 30011 strain) was obtained from the American Type Culture Collection. Following established protocols, trophozoites were cultured axenically in peptone-yeast-glucose (PYG) medium at 26°C.

Caco-2 cells were were cultured in MEM medium (Corning, USA) with 20% (v/v) fetal bovine serum (HyClone, Australia); HBEC-5i cells were cultured in DMEM/F12 medium (Corning, USA) with 10% (v/v) fetal bovine serum and growth factor supplements (Corning, USA); HCEC, and HEP-2 cells were cultured in DMEM (Corning, USA), with 10% (v/v) fetal bovine serum. All the cells were cultured with penicillin–streptomycin (100 U/mL; Gibco) at 37°C in a 5% CO2 incubator.

### Compounds

Isoliquiritigenin, liquiritigenin, liquiritin, licochalcone A, licochalcone B, licochalcone E, dipotassium glycyrrhizinate, glabridin, glycyrrhizic acid, and ammonium glycyrrhizinate were obtained from MCE (MedChem Express, Monmouth Junction, NJ, USA). These compounds were dissolved in 100% dimethyl sulfoxide (DMSO; Sigma-Aldrich, St. Louis, MO, USA) and further diluted in PYG medium to achieve the desired experimental concentrations for *in vitro* experiments.

### Cell viability assay and IC_50_ determination

*A. castellanii* trophozoites (1 × 10^4^ cells per well) were seeded in a 96-well white microplate (Eppendorf, Hamburg, Germany) and incubated with a medium containing varying drug concentrations at 26°C for 24 h, 48 h and 72 h, after which morphological changes were observed under an inverted microscope (Olympus). Cell viability assay results were assessed employing CellTiter-Glo (Promega, Madison, WI, USA), according to the guidelines of the manufacturer. Luminescence was recorded on a modular multimode microplate reader (BioTek Synergy H1), with IC_50_ values of the compound and growth curves subsequently generated utilizing GraphPad Prism 8. Each experiment was conducted independently in triplicate.

### Cytotoxicity assay in HCEC and Caco-2 cells

HCEC and Caco-2 cells were cultured at a density of 1 × 10^4^ cells per well in a 96-well plate and allowed to adhere overnight. The cells were then treated with serially diluted ISL and GLA at final concentrations ranging from 0 to 160 μM or 0 to 64 μM, respectively, and incubated for 24 h. Cell viability was evaluated using a Cell Counting Kit (CCK-8) reagent (Dojindo, cat.CK04), following the instructions of the manufacturer. Absorbance was measured at 450 nm following 1-h incubation with the reagent. The CC_50_ was determined employing GraphPad Prism 9.1.0 software (GraphPad Software).

### Hoechst33342/PI double stain assay

To investigate the mode of cell death, PI and Hoechst 33342 fluorescent staining (62249; Thermo Scientific™) were performed employing a confocal microscope. Trophozoites were cultured in 48-well plates and treated with 0.1% DMSO, ISL, and GLA for 24 h. After washing twice with PBS, the trophozoites (5×10^5^/mL) were then incubated with Hoechst 33342 (25 μg/mL) and PI (10 μg/mL) at 26°C for 10 min in the dark. Fluorescence images were then captured using a Leica TCS SP8 microscope.

### Caspase-3 activity assay

Caspase-3 activity was detected employing NucView® 488 Caspase-3 Assay Kit (Biotium, Inc., Fremont, CA, USA). Following 24 h of ISL and GLA treatment, trophozoites (1 × 10^5^ cells/well) were incubated with 100 μL of 5 μM NucView® 488 substrate at room temperature for 30 min in the dark. Cells were observed directly in the medium containing the substrate. Fluorescence intensity (λEx = 485 nm and λEm = 515 nm) was measured utilizing a modular multimode microplate reader, and fluorescence images were captured using a Leica TCS SP8 microscope.

### Flow cytometry analysis

*A. castellanii* trophozoites (2 × 10^6^ cells/flask) were treated with various concentrations of ISL or GLA for 24 h. The ROS level in trophozoites was assessed utilizing a ROS assay kit (R252, Dojindo Molecular Technologies, Inc., Kumamoto, Japan), following the instructions from the manufacturer. Fluorescence-activated cell sorting (FACS) was performed on a FACSAria instrument (BD Biosciences, San Jose, CA, USA) with a 488-nm argon excitation laser. The analysis gates were defined utilizing untreated amoebae, and data analysis was conducted using FlowJo 10.8.1 software (FlowJo LLC, Ashland, OR, USA).

### *Acanthamoeba-*mediated cell cytotoxicity assays

Cytopathogenicity assays were performed to assess *Acanthamoeba*-mediated human cell damage. Briefly, *A. castellanii* trophozoite was treated with ISL (55 μM and 75 μM) or GLA (8 μM and 16 μM) for 180 min at 26□. Trophozoites treated with 0.1% DMSO served as control. The cultures, including pre-treated trophozoites, were washed twice with an FBS-free MEM medium and resuspended in the corresponding FBS-free cell culture medium. The trophozoites were then added to HBEC-5i, HCEC, or HEP-2 cells at a ratio of 1:5 in either 96-well plates (for CCK-8 assay) or 24-well plates (for qPCR assay). The co-cultures were incubated for 24 h in a cell culture incubator at 37□ in a 5% CO_2_. The *Acanthamoeba*-mediated cell death was indirectly quantified using CCK-8 reagent (DOJINDO, Japan). Following three distinct, independent trials, the data are presented as the mean ± standard error.

### Mitochondrial membrane potential and mitochondrial superoxide assay

The mitochondrial membrane potential (MMP) and mitochondrial superoxide (mtSOX) levels were evaluated utilizing the JC-1 MitoMP Detection Kit (MT09, Dojindo) and mtSOX Deep Red Detection Kit (MT14, Dojindo). After 24 h of ISL or GLA treatment, JC-1 solution was added to the culture medium to a final concentration of 4 μM. Subsequently, treated trophozoites (1×10^4^ per tube) were incubated at 26°C for 30 min. After washing twice with HBSS, the trophozoites were then incubated in 10 μM mtSOX Deep Red working solution at 26°C for 30 min. The fluorescence (Green: λEx = 488 nm and λEm = 528 nm, Red: λEx = 561 nm and λEm = 600 nm, Deep red: λEx = 550 nm and λEm = 675 nm) was measured using a modular multimode microplate reader (BioTek Synergy H1) and the fluorescence images were obtained employing a Leica TCS SP8 microscope.

### Measurement of the superoxide dismutase activity

Trophozoites (1 × 10^7^) were collected from each group and were washed twice with an equal volume of PBS at 800 g for 10 min. The suspension was ultrasonicated (80 W, 1 s intervals, 10 min) and then centrifuged at 10,000 ×g for 15 min. The SOD activity in each fraction was measured utilising a SOD Assay Kit-WST (Dojindo Molecular Technologies, Tokyo, Japan). After incubating the plate at 37°C for 20 min, absorbance at 450 nm was measured, and the inhibition rate was calculated following the instructions from the manufacturer.

### NAD^+^/NADH ratio assay

Trophozoites (5 × 10^6^ / mL) were collected from the following DMSO-treated groups: 55 μM ISL-treated group and 75 μM ISL-treated groups. Samples were washed twice with an equal volume of PBS at 800 g for 10 min. After adding 1 mL of PBS, the samples were sonicated (80 W, 1 s intervals, 5 min). The cellular NAD^+^/NADH ratio was quantified using an NAD^+^/NADH Assay Kit-WST (Dojindo Molecular Technologies, Tokyo, Japan), according to the instructions of the manufacturer. After incubating the plate at 37°C for 1 h, the absorbance was measured at 450 nm.

### 7-DHC detection assay

Trophozoites (1 ×10^7^ / mL) were collected from the DMSO-treated group: 8 and 16 μM GLA-treated groups. The samples were washed twice with an equal volume of PBS at 800 g for 10 min. After adding 1 mL of PBS, the samples were sonicated (80 W, 1 s intervals, 5 min). Cellular 7-Dehydrocholesterol (7-DHC) was quantified utilizing a 7-DHC ELISA kit (DLdevelop, Jiangsu, China), which is a competitive inhibition enzyme immunoassay technique. Absorbance at 450 nm was measured utilizing a modular multimode microplate reader (BioTek Synergy H1).

### qPCR analysis

Total RNA was isolated from *A. castellanii* trophozoites and host cells as previously described (Chen et al., 2024). The reactions were performed in a 96-well plate using SYBR Premix Ex Taq (Takara, Dalian, Liaoning, China), with primers targeting *A. castellanii PARP*, *NAMPT*, *NMNAT*, *GST*, *Sir2*, *NQO1*, *SC5D*, *ACAT*, *CYP51*, *ERG4/ERG24*, *SC4MO,* and *ERG5-like*. qPCR was also conducted to measure the expression of apoptosis-related genes, including *APAF-1*, *ATM*, *BCL-2*, *BAX*, *TNF-*α, *TGF-*β, *FAS,* and *SOD2* in the host cells. Primers for these genes, sourced from published sequences (**Table S1**), were utilized for qPCR analysis on an ABI 7500 RT-PCR system (Applied Biosystems, Foster City, CA, USA). The relative expression of each primer between the control and treatment groups was calculated using the 2^-△△Ct^ method, with values normalised to the *18S* reference housekeeping gene.

### Transcriptome data analysis

The trophozoites were treated with 55 μM ISL or 8 μM GLA for 24 h. Total RNA was then isolated from the trophozoites using the RNAmini kit (Qiagen, Germany). Strand-specific RNA libraries were prepared with the TruSeq RNA sample preparation kit (Illumina, San Diego, CA, USA). Sequencing was then performed on the Illumina Novaseq 6000 platform. The raw data were subsequently analysed as previously described (Dong et al., 2024). The protein-protein interaction network was visualised using the STRING database and analysed with k-means clustering to identify the hub genes (Szklarczyk et al., 2019).

## Statistical analysis

The data were presented as mean ± standard deviation (SD) and were analyzed using a one-way ANOVA test. All data were visualised using GraphPad Prism, and *p* < 0.05 was regarded as statistically significant.

## Results

### Screening of ISL and GLA from licorice derivatives that could suppress viability of *A. castellanii*

To evaluate the *in vitro* efficacy of licorice derivatives in *A. castellanii*, we selected 10 drugs for testing using the CellTiter-Glo® Luminescent cell viability assay. Compared to that of the DMSO-treated group, ISL, LicoA, and GLA induced a significant dose-dependent decrease in the trophozoite viability (**Fig. 1A and 1B**). Additionally, the viability assay revealed that ISL and GLA induced a marked reduction time-dependently (**Fig. 1C**). The IC_50_ values of ISL and GLA were estimated to be 53.71±2.04 μM and 8.59±0.43 μM, respectively, after 24 h treatment. Exposure to ISL and GLA at IC_50_ did not impair the viability of HCEC and Caco-2 cell lines, while LicoA exhibited significant cytotoxicity (**Fig. 1C; Fig. S1**). The data represent means ± SDs from three independent experiments. Simultaneously, trophozoites progressively became rounded and elliptical, with spinous protrusions and the disappearance of intracellular vacuoles, showing signs of encystment after 24 h of treatment (**Fig. 1D**). Overall, ISL and GLA exhibited more potent therapeutic efficacy compared to those of other selected licorice derivatives. Therefore, we selected ISL concentrations of 55 and 75 μM and GLA concentrations of 8 and 16 μM for the subsequent experiments.

**Fig. 1.**
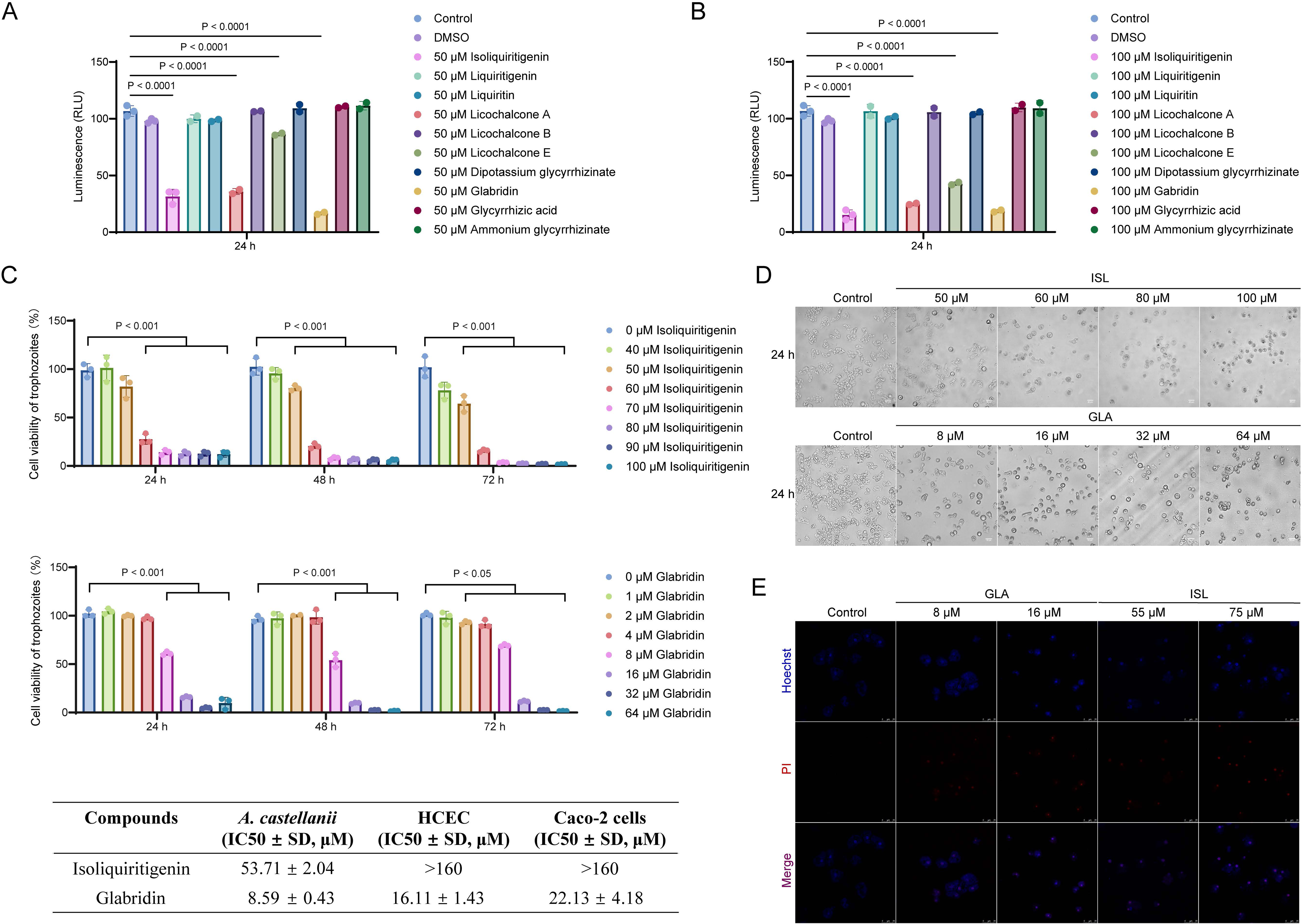
Suppressing *A. castellanii* trophozoites viability by isoliquiritigenin (ISL) and glabridin (GLA). (A, B) Viability of trophozoites following 50 μM and 100 μM licorice derivatives treatments for 24 h determined using the CellTiter-Glo assay. (C, D) Viability of trophozoites following 40-100 μM ISL and 1-64 μM GLA treatments for 24, 48, and 72 h. (E) IC_50_ values of ISL and GLA in *A. castellanii*, HCEC and Caco-2 cells. Data are mean ± SD of n = 3 biological replicates. (F) Morphological changes on trophozoites incubated with control (0.1% DMSO) and different concentration of ISL and GLA for 24 h. Scale bar: 20 μm. (G) Images of trophozoites after 24 h of ISL and GLA treatment with Hoechst 33342 - PI staining. Scale bar: 25 μm.

### ISL and GLA trigger apoptotic death of trophozoites and increase intracellular oxidative stress damage

To determine whether the inhibitory effect of ISL and GLA on cell viability is associated with apoptosis induction, we assessed the mode of cell death after 24 h of culture. The cells were stained with Hoechst (blue) and PI (Red) to visualise the nuclei of live or dead cells, respectively. Hoechst 33342 emitted blue fluorescence, which indicates nuclear alterations that distinguish apoptotic cells from necrotic and viable cells, while the red fluorescence emitted by PI indicates DNA strand damage and marks dead cells within a population. Compared to the control group, the condensation of chromatin in *A. castellanii* trophozoites treated with ISL and GLA, especially at the high concentration of GLA, exhibited strong blue and red fluorescence (**Fig. 1E**). Activation of caspase-3 is recognised as a key event in the induction of apoptosis. Consistent with above results, caspase-3 production was also increased in *A. castellanii* treated with ISL and GLA (**Fig. 2A**). Strong green fluorescence signals were observed at nuclear sites in the 75 μM ISL-treated and 16 μM GLA-treated groups, while a smaller amount of green fluorescence signal was observed in the 55 μM ISL-treated and 8 μM GLA-treated groups (**Fig. 2B**). No significant fluorescence signals were observed in the control groups (**Fig. 2B**). The findigs suggest that ISL and GLA significantly induced apoptosis in *A. castellanii* trophozoites at the corresponding concentrations.

**Fig. 2.**
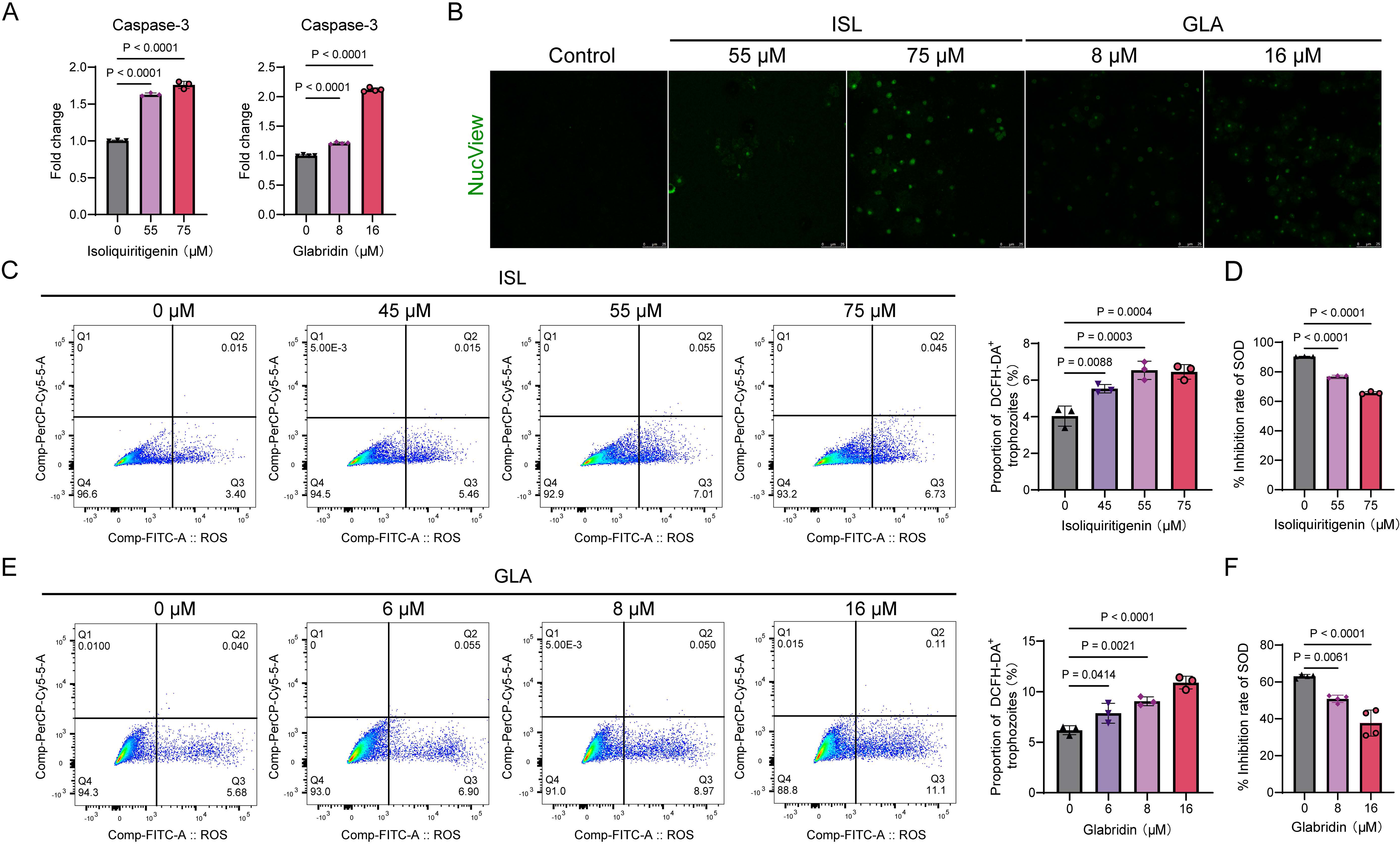
ISL and GLA increased the level of caspase-3 and ROS and decreased the expression of SOD (A) Caspase-3 activity of trophozoites incubated with control and different concentration of ISL and GLA for 24 h. (B) Images of trophozoites after 24 h of ISL and GLA treatment with NucView staining. (C) ROS level after 24 h of ISL was measured using DCFH-DA flow cytometry. (D) Inhibition rate of SOD after 24 h of ISL treatment. (E) ROS level after 24 h of GLA was measured using DCFH-DA flow cytometry. (F) Inhibition rate of SOD after 24 h of GLA treatment.

To further investigate the inherent condition of trophozoites, we evaluated intracellular ROS production using flow cytometry. The fluorescent indicator DCFH-DA was employed to evaluate the total ROS production. **Fig. 2C and 2E** illustrate that the ISL-treated and GLA-treated groups exhibited an increase in intracellular ROS levels dose-dependently. Additionally, ISL and GLA reduced the expression of SOD (**Fig. 2D and 2F**). This served as direct evidence and theoretical support that ISL and GLA significantly enhanced ROS generation and weakened cellular antioxidant defense mechanisms, leading to an imbalance in redox status.

### ISL and GLA disrupt mitochondrial homeostasis in *A. castellanii* trophozoites

Mitochondria are dynamic organelles essential for ATP production (22). Here, we explored whether ISL or GLA influences mitochondrial function and integrity. To assess the changes in mitochondrial dynamics, we analysed the alteration in mitochondrial membrane potential using the JC-1 probe and measured mitochondrial superoxide production using mtSOX dyes. The results showed that the red/green fluorescence ratio of trophozoites in the ISL-treated and GLA-treated groups was significantly lower than those of the control group, indicating a disruption of mitochondrial membrane potential (**Fig. 3A and 3B**). Additionally, both ISL and GLA increased mitochondrial ROS levels (**Fig. 3A and 3C**). Elevated mitochondrial ROS generation damages mitochondrial components, leading to dysfunctional mitochondrial accumulation. These findings confirm that ISL and GLA induce mitochondrial damage in *A. castellanii* trophozoites.

**Fig. 3.**
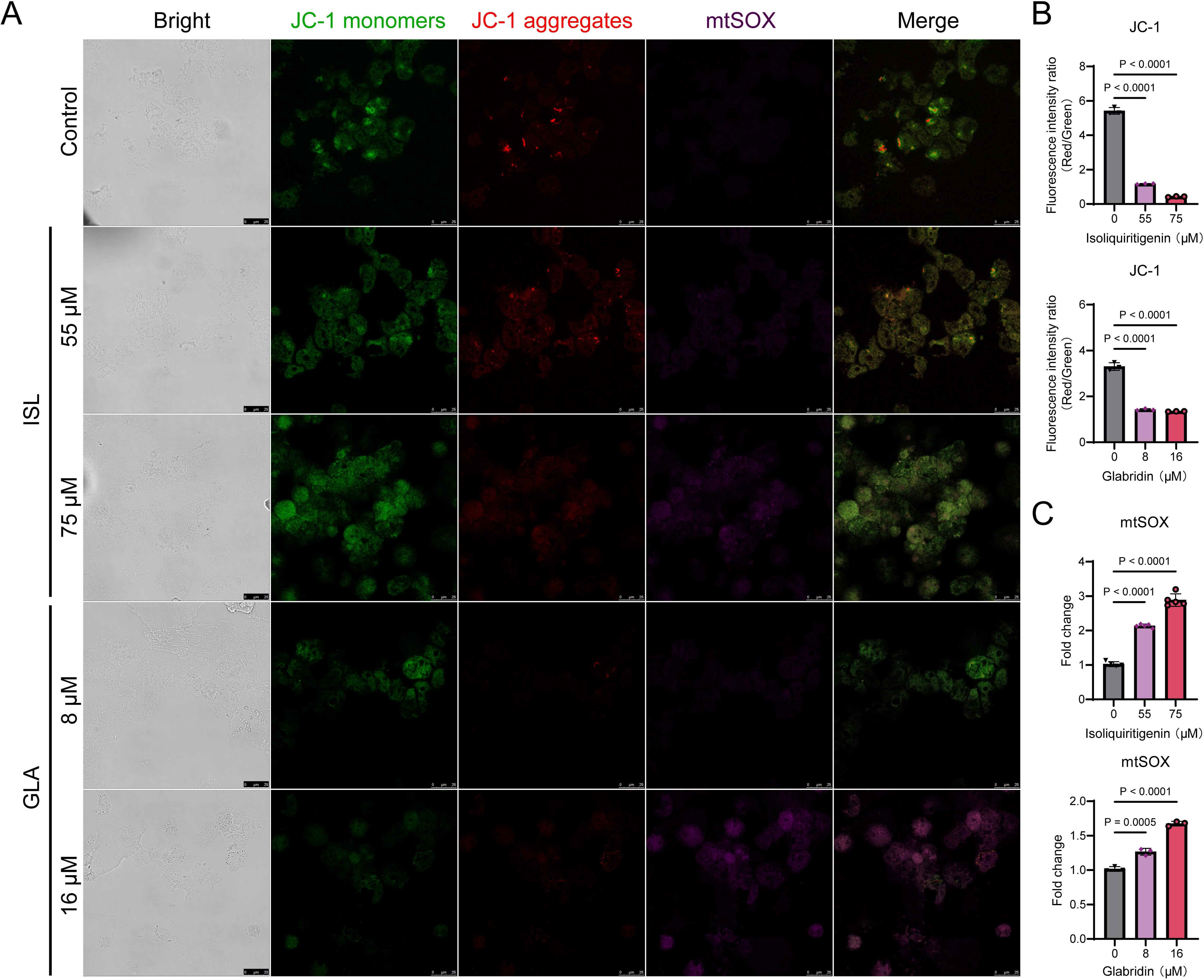
ISL and GLA disrupt mitochondrial homeostasis in *A. castellanii* (A) Images of trophozoites after 24 h of ISL and GLA treatment with JC-1 and mtSOX staining. Scale bar: 25 μm. (B) Fluorescence intensity ratio (Red/Green) of trophozoites after 24 h of ISL and GLA treatment. (C) Mitochondrial ROS levels of trophozoites after 24 h of ISL and GLA treatment.

### ISL and GLA effectively prevent host cells from *A. castellanii* trophozoites invasion

The findings above suggest that ISL and GLA effectively inhibit amoebic activity. To explore whether ISL and GLA could protect host cells from trophozoite invasion, cell invasion assays were conducted. The CCK-8 results indicated that ISL and GLA at IC_50_ doses effectively inhibited trophozoite invasion into the three host cell lines, with ISL showing near-complete protection of the host cells (**Fig. 4A**). In the RT-qPCR analysis, the mRNA expression levels of *FAS, SOD2, BAX,* and *TNF-*α were down-regulated, while mRNA expression levels of *ATM,* and *BCL2* were up-regulated in ISL or GLA treated groups (**Fig. 4B**). The results also demonstrated that amoebae invasion increased the expression of multiple apoptosis-related, cytokine, and antioxidant genes in the host. However, in the ISL and GLA pre-treated groups, the expression of these genes decreased to varying degrees. These findings suggest that ISL and GLA can effectively prevent host cell invasion by *A. castellanii* trophozoites.

**Fig. 4.**
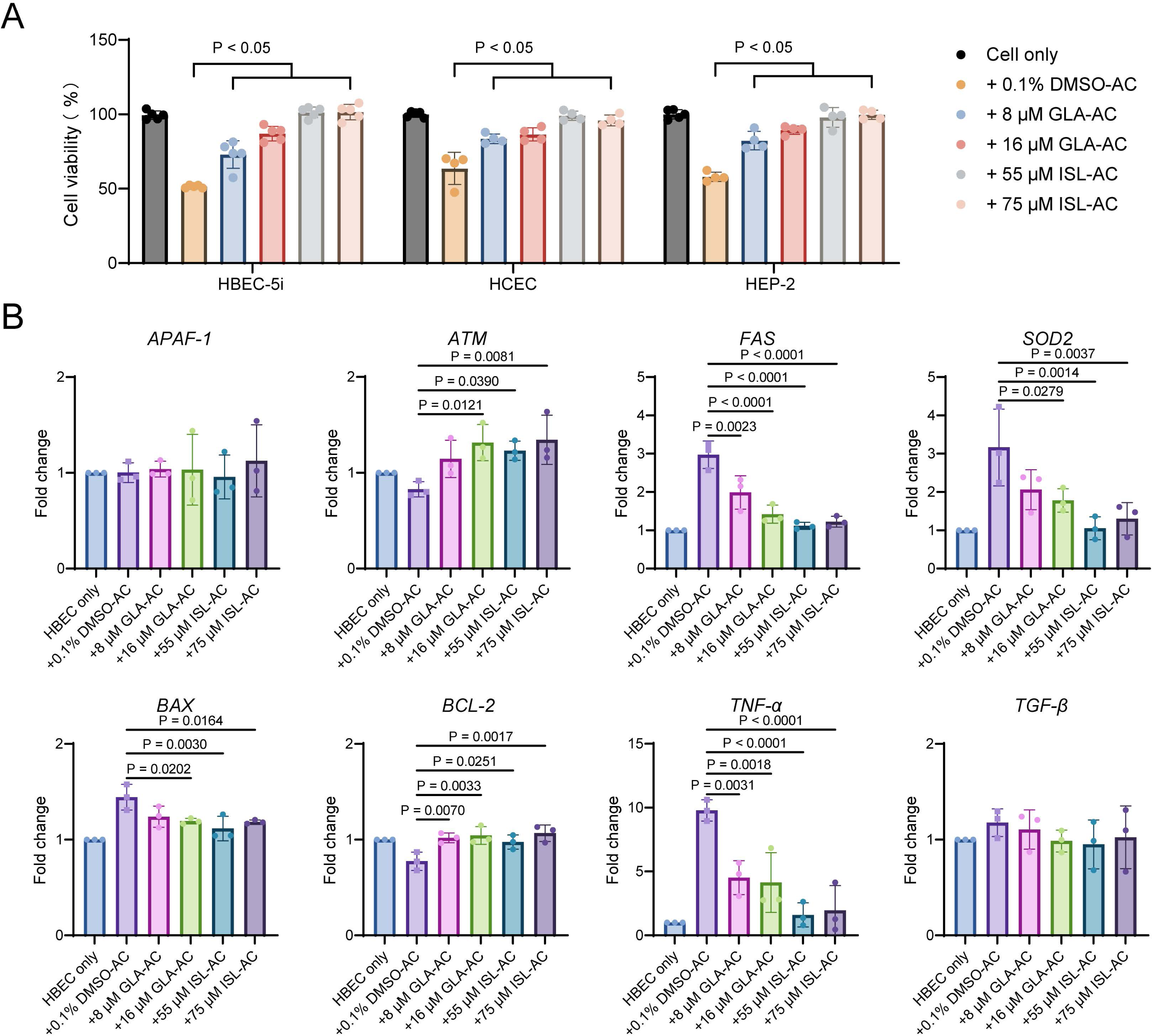
ISL and GLA effectively prevent host cells from *A. castellanii* trophozoites invasion (A) The CCK-8 results of three host cell lines cocultured with pre-treated trophozoites for 24 h. (B) Relative mRNA expression of *APAF-1*, *ATM, FAS, SOD2, BAX, BCL-2, TNF-*α*, and TGF-*β under pre-treated trophozoite treatments for 24 h in HBEC-5i cells. Gene expression was normalised to *GAPDH* expression levels.

### ISL affects the NAMPT/ NAD^+^/SIRT pathway in *A. castellanii*

RNA sequencing was performed to gain insight into the potential protective effects of ISL and GLA on host cells against amoebic invasion, alongside the molecular mechanisms underlying *A. castellanii* suppression.

Trophozoites of *A. castellanii* were treated with ISL or GLA at IC_50_ doses and then subjected to RNA sequencing to analyse alteration in intracellular gene expression profiles. After 24 h of ISL treatment, 495 genes exhibited significant changes, with 359 up-regulated and 136 downregulated [log2FC < −1 or log2FC > 1; *p* < 0.05] (**Fig. 5A)**. Gene Ontology (GO) enrichment analysis of these differentially expressed genes revealed that ISL primarily affected the NAD^+^ biosynthetic process, and nicotinate-nucleotide adenylytransferase activity (**Fig. 5B**). GO network diagram indicated that ISL might influence not only the NAD^+^ biosynthetic process but also intracellular signal transduction (**Fig. 5C**). The PPI network analysis identified 16 key proteins involved in nicotinate and nicotinamide metabolism. Among these, the expression levels of nine genes were significantly up-regulated (e.g., nicotinamide-nucleotide adenylytransferase and transcriptional regulator sir2 family protein), while three genes were significantly downregulated (*p* < 0.05) (**Fig. S2**). Additionally, Sankey diagrams of KEGG enrichment provided a more intuitive visualisation, highlighting two genes, *NMNAT* and *NAMPT*, associated with the nicotinate and nicotinamide metabolism pathway (**Fig. 5D**). The volcano plot showed that the expression of *NNMT* was down-regulated (**Fig. 5A**). Consistent with the RNA sequencing results, the mRNA expression levels of *NMNAT*, *NAMPT*, *sir2*, and *PARP* were up-regulated after both 55 μM and 75 μM ISL treatments (**Fig. 5E**), all of which play crucial roles in the NAMPT/NAD^+^/SIRT pathway (**Fig. 5F**). The mRNA expression levels of *GST* and *NQO1* were also up-regulated (**Fig. 5E**). Furthermore, the biochemical assay showed that ISL significantly decreased NAD^+^/NADH ratio (**Fig. 5G**). These findings suggest that ISL may exert its effects by modulating the NAMPT/NAD^+^/SIRT pathway in *A. castellanii* trophozoites, thereby inducing oxidative stress damage to the trophozoite.

**Fig. 5.**
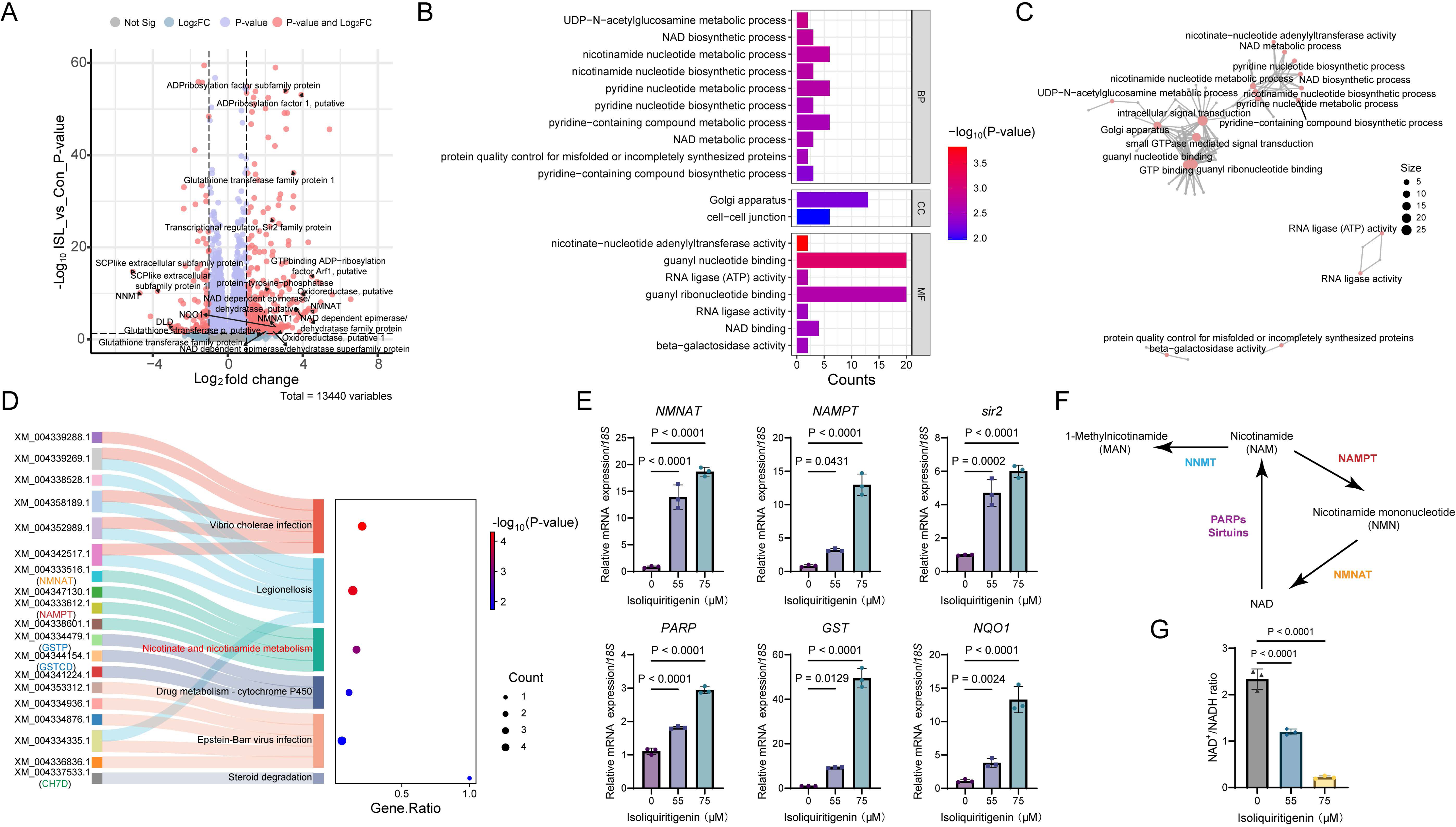
ISL affected the NAD metabolic process in *A. castellanii* trophozoites (A) Volcano plot of the DEGs in trophozoites under 55 μM ISL treatments. (B) GO enrichment analysis filtered based on a P-value ≤ 0.05 in trophozoites treated with ISL. BP: biological process. CC: cellular component. MF: molecular function. (C) A network diagram of the top 20 GO terms and candidate genes, ranked by-Log_10_(P-value), is displayed. The node size represents the total number of candidate genes associated with each GO term. (D) Sankey diagrams of KEGG enrichment (E) Relative mRNA expression of *NMNAT*, *NAMPT*, *sir2*, *PARP*, *GST* and *NQO1* under 55 μM ISL treatments for 24 h in *A. castellanii* trophozoites. Gene expression was normalised to *18S* expression levels. (F) Schema diagram of NAMPT/ NAD^+^/SIRT pathway (G) NAD^+^/NADH ratio of trophozoites after 24 h of 55 μM and 75 μM ISL treatment.

### GLA might impact the sterol metabolic process in *A. castellanii*

After 24 h of GLA-treatment, 607 genes exhibited significantly alteration, with 340 up-regulated and 267 down-regulated [log2FC < −1 or log2FC > 1; p < 0.05] **(Fig. 6A**). The volcano plot revealed that the expression of acetyl-CoA cholesterol acyltransferase (ACAT) and cytochrome b5 domain-containing protein were down-regulated, while one lipoprotein gene (UniProt IDs: L8H5W8) was up-regulated (**Fig. 5A**). GO enrichments analysis identified the ergosterol, phytosteroids, and cellular lipid biosynthetic processes among top-ranked pathways (**Fig. 6B**). The GO network diagram illustrated the interactions between acetyl-CoA acyltransferase activity, cellular lipid biosynthesis, and ergosterol biosynthesis (**Fig. 6C**). The PPI network revealed that among the 36 key proteins involved in the steroid metabolic pathway, the expression levels of 17 genes were significantly down-regulated (e.g., C4-methyl sterol oxidase (SC4MO) and ergosterol biosynthesis ERG4/ERG24 family protein), while 4 genes were significantly up-regulated (e.g., mevalonate kinase and cytochrome P450 superfamily proteins) (*p* <0.05) (**Fig. S3**). Sankey diagram based on down-regulated KEGG terms illustrated the gene *SC5D* and *CH7D* (**Fig. 6D**), which are the vital enzymes to synthesize 7-DHC. In our qPCR analysis, the mRNA expression levels of *SC5D*, *ACAT*, *CYP51*, *ERG4/ERG24*, *SC4MO* and *ERG5-like* were down-regulated both in 8 μM and 16 μM GLA-treated group (**Fig. 6E**). The ELISA assay also showed that GLA significantly decreased 7-DHC level (**Fig. 6F**). These results highlight the potential role of GLA in influencing *A. castellanii* trophozoites via modulation of the sterol metabolism pathway.

**Fig. 6.**
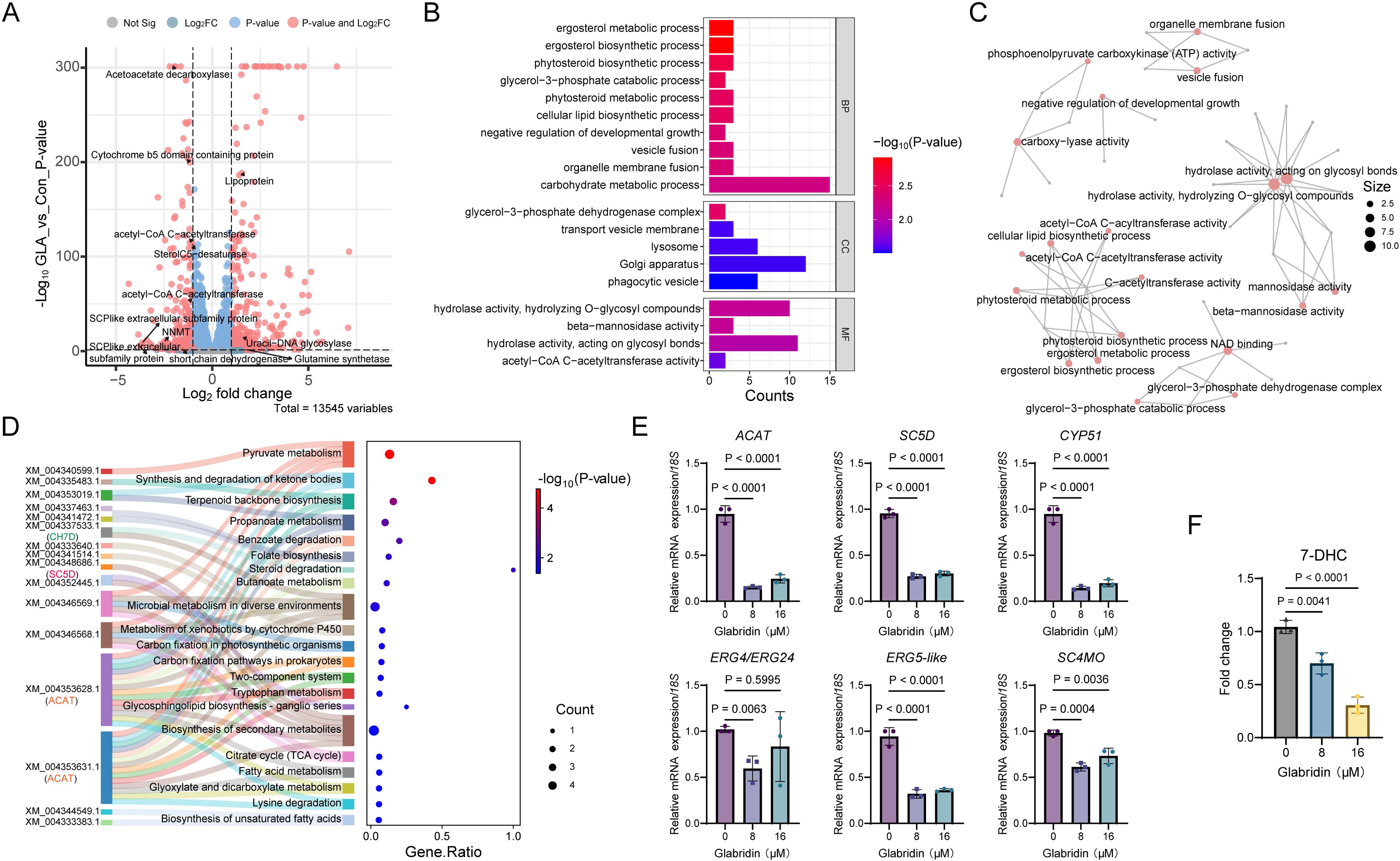
GLA impacted the sterol metabolic process in *A. castellanii* trophozoites (A) Volcano plot of the DEGs in trophozoites under 8 μM GLA treatments. (B) GO enrichment analysis filtered based on a P-value ≤ 0.05 in trophozoites treated with GLA. BP: biological process. CC: cellular component. MF: molecular function. (C) A network diagram of the top 20 GO terms and candidate genes, ranked by-Log_10_(P-value), is displayed. The node size represents the total number of candidate genes associated with each GO term. (D) Sankey diagrams of down-regulated KEGG enrichment (E) Relative mRNA expression of *SC5D* and *DHCR7* under 8 μM GLA treatments for 24 h in *A. castellanii* trophozoites. Gene expression was normalised to *18s* expression levels. (F) 7-DHC level of trophozoites after 24 h of 8 μM and 16 μM GLA treatment.

## Discussion

Numerous research has explored the use of natural remedies to combat tumours, bacterial infections, and parasitic diseases (Chen et al., 1994; Hsieh et al., 2016; Lê et al., 2023; Wu et al., 2024). The therapeutic potential of *G. glabra—* a common perennial herb with centuries of use in traditional Chinese medicine—has attracted significant attention. Nevertheless, its effects and mechanisms in inhibiting *Acanthamoeba* activity have not been extensively investigated.

In this study, 10 licorice extracts were selected to evaluate their anti-*Acanthamoeba* activity. The findings provide evidence that ISL and GLA effectively suppressed the growth of *A. castellanii in vitro* dose-and time-dependently. Furthermore, exposure to ISL and GLA at their IC_50_ did not influence the normal viability of HCEC and Caco-2 cell lines. Recently, the safety of ISL and GLA as drug candidates has been experimentally confirmed. Further, the *in-vitro* cytotoxicity of ISL was validated using a HEK cell line, revealing that ISL was non-toxic at concentrations up to 100 μg/mL (Kumar, 2024). While Lico A also exhibited an inhibitory effect, it showed greater cytotoxicity.

Microscopic observation revealed that ISL and GLA induced a concentration-dependent decline in the number of trophozoites and significant morphological alterations. To determine whether drug-induced cell death was associated with apoptosis, we utilised Hoechst/PI double staining to examine nuclear morphological changes and identify apoptotic cells. Furthermore, caspase-3 activity was measured to provide further evidence of its involvement in the observed cell death. Chromatin condensation in *A. castellanii* trophozoites treated with ISL and GLA was clearly visible, as indicated by intense blue and red fluorescence. Consistent with this result, caspase-3 production was also increased in *A. castellanii* treated with ISL and GLA, suggesting apoptotic changes in the trophozoites. Multiple pieces of evidence suggest that an increase in ROS production or the disruption of cellular redox balance can induce cell cycle arrest or apoptosis by regulating DNA damage and metabolic processes. Our flow cytometric analysis revealed that ISL and GLA significantly increased cellular ROS production dose-dependently. The levels of SOD were significantly decreased, as demonstrated by experimental results. Large amounts of ROS may be released into the cytoplasm owing to electron leakage from the mitochondrial electron transport chain during mitochondrial fragmentation (Nunnari and Suomalainen, 2012; Areti et al., 2016). Our findings suggest that ISL and GLA may lead to mitochondrial dysfunction, as confirmed by the significantly increased mitochondrial ROS generation and disruption of mitochondrial membrane potential.

Based on the transcriptomic signature and functional properties of ISL-treated trophozoites, nicotinate and nicotinamide metabolism were significantly clustered in both GO enrichment and KEGG enrichment analyses. NAD^+^ is a crucial metabolite necessary for sustaining cellular energy metabolism and exists in two interconvertible forms: NAD^+^ and its reduced counterpart, NADH (Migaud et al., 2024). Additionally, NAD^+^ acts as a second messenger involved in numerous cellular processes, including mitochondrial function, anti-oxidation defense, gene expression, and cell death. Consistent with the RNA sequencing results, the mRNA expression levels of *NMNAT*, *NAMPT, sir2*, and *PARP* were up-regulated after ISL treatment. NAMPT is the rate-limiting enzyme in a key NAD^+^ salvage pathway, while PARPs and SIRTs are NAD^+^ degrading enzymes, both of which maintain sufficient levels for cell survival (Fulco et al., 2008; Sampath et al., 2015). AcSir2 (ACA1_084880) was classified as a class□IV sirtuin and exhibited functional SIRT deacetylase activity (Joo et al., 2020). The discovery suggests that yeast Sir2 and its homologues might function as nutrient and metabolic sensors, relaying nuclear changes in the NAD^+^/NADH ratio to regulate transcription and genome stability (Bonkowski and Sinclair, 2016). Biochemical assay further demonstrated that ISL significantly reduced the NAD^+^/NADH ratio, suggesting that ISL may disrupt the NAD^+^ homeostasis in trophozoites. While the Nrf2 signalling pathway in *A. castellanii* remains unclear, its downstream effector enzyme *NQO1* expression was up-regulated under ISL conditions. In conclusion, these findings suggest that ISL may influence NAD^+^/PARPs/SIRTs signalling transduction in *A. castellanii* trophozoites. However, the current results are insufficient to draw definitive conclusions, indicating that further research is needed.

Transcriptomic analysis of GLA revealed that GO and KEGG pathway enrichment analyses indicated the modulation of sterol metabolism, including processes such as ergosterol metabolic process, acetyl-CoA C-acyltransferase activity, and steroid degradation. A previous study suggests that GLA might disrupt sterol metabolism in *F. graminearum* by modulating ergosterol biosynthesis proteins, including ERG4/ERG24, and reducing ergosterol levels (Yang, 2021). Ergosterol, a key component of fungal and parasitic membranes, is essential for maintaining membrane integrity, and its depletion induces membrane leakage and cell death, highlighting its potential as a drug target (Scalese, 2024; Shing et al., 2020). A recent study demonstrates that ergosterol and 7-DHC, both containing an indistinguishable B-ring structure, affect oxidative stress levels and regulate cell sensitivity to ferroptosis (Freitas et al., 2024). In mammalian cholesterol biosynthesis, 7-DHC is synthesised by SC5D and converted to cholesterol by DHCR7. In *C. elegans*, this process is reversed, with CH7D catalysing the conversion of cholesterol to 7-DHC (Li et al., 2024; Wollam et al., 2011). Isotopic labelling in *N. fowleri* revealed enrichment of ^13^C in 7-DHC, indicating that this metabolite originates from *de novo* biosynthesis and the metabolism of scavenged cholesterol (Zhou et al., 2018a). Our findings suggest that two genes involved in this biosynthesis, *SC5D* and *CH7D*, were significantly down-regulated after GLA treatment. Consistently, our assay confirmed that GLA treatment significantly decreased the levels of 7-DHC. Additionally, we observed that several key genes involved in the sterol biosynthetic pathway, including *ACAT*, *CYP51*, *ERG4/ERG24*, *SC4MO,* and *ERG5-like*, were significantly down-regulated. CYP51 is a well-known target for azole antifungal and amoebicidal effects (Lamb et al., 2015; Monk and Keniya, 2021; Sharma et al., 2023). Studies demonstrate that AcCYP51 preferentially utilises obtusifoliol as a substrate over lanosterol (Hargrove et al., 2024; Zhou et al., 2018b). Nevertheless, the specific role of ACAT, ERG4/ERG24, SC4MO, and ERG5-like proteins in *A. castellanii* remains unclear. In humans, ACAT is the primary enzyme responsible for the intracellular esterification of free cholesterol, catalysing its reaction with fatty acyl-CoA to form cholesterol esters (Markina et al., 2022). The sterol C4-methyl oxidase plays a role in the sterol C4-demethylation process in *Leishmania* (Jin et al., 2024). Additionally, overexpression of the CYP710C1 gene, which encodes sterol C-22 desaturase, confers resistance to amphotericin B in *L. donovani* (Bansal et al., 2019). Further investigations are required to elucidate their precise roles in *A. castellanii*.

Our research suggests that the bioactive compounds in licorice, ISL, and GLA may disrupt key physiological processes in *A. castellanii*, resulting in mitochondrial dysfunction, increased oxidative stress, and reduced virulence toward host cells. This evidence highlights the potential of combining traditional herbal remedies with modern treatments, providing a promising alternative or supplement to existing anti-parasitic medications.

## Conclusion

Our study demonstrated that ISL and GLA suppressed the viability of *A. castellanii* trophozoites and induced apoptosis. Both ISL and GLA increased cellular oxidative stress and disrupted mitochondrial homeostasis in trophozoites. Further analysis revealed that the anti-*Acanthamoeba* activity of ISL on trophozoites may be associated with the regulation of NAD metabolism, while the inhibitory effect of GLA may be partially owing to its effect on sterol metabolism. These findings suggest that ISL and GLA exhibit strong anti-*Acanthamoeba* activity with low cytotoxic effects *in vitro*, making them promising candidates for developing much-needed anti-*Acanthamoeba* drugs.

## Abbreviations

ACAT: acetyl-CoA cholesterol acyltransferase
AK: acanthamoeba keratitis
APAF-1: apoptotic protease activating factor-1
ATM: ataxia telangiectasia-mutated gene
CC_50_: 50% cytotoxicity concentration
CH7D: cholesterol 7-desaturase
CYP51: sterol 14α-demethylase
DCFH-DA: 2′,7′-dichlorodihydrofluorescein diacetate
DMSO: dimethyl sulfoxide
ERG4/ERG24: ergosterol biosynthesis ERG4/ERG24 family protein
ERG5-like: Sterol C22 desaturase-like
FACS: fluorescence-activated cell sorting
GLA: glabridin
GO: gene Ontology
GST: glutathione transferase
IC_50_: 50% inhibitory concentration
ISL: Isoliquiritigenin
Lico A: licochalcone A
MDA: malondialdehyde
MMP: mitochondrial membrane potential
NAD^+^: nicotinamide adenine dinucleotide
NAMPT: nicotinamide phosphoribosyltransferase
NNMT: nicotinamide N-methyltransferase
NMNAT: nicotinamide-nucleotide adenylyltransferase
NO: nitric oxide
NQO1: NADPH: quinone reductase
PARP: poly ADP-ribose polymerase
PI: propidium iodide
PYG: peptone-yeast-glucose
ROS: reactive oxygen species
SC4MO: C4-methyl sterol oxidase
SC5D: sterol C5-desaturase
SD: standard deviation
SOD: superoxide dismutase
TNF-α: tumour necrosis factor-alpha
TGF-β: transforming growth factor beta
7-DHC: 7-dehydrocholesterol.

## Data availability statement

The raw data of NGS will be submitted to NCBI and publicly disclosed before publication.

## Author contributions: CRediT

**Lijun Chen**: Conceptualization, Data curation, Formal analysis, Methodology, Software, Writing – original draft. **Meng Feng**: Investigation, Methodology, Visualization. **Dai Dong**: Data curation, Methodology. **Wenwen Jing**: Resources, Software, Supervision. **Qingtong Zhou**: Software, Supervision. **Xunjia Cheng**: Conceptualization, Funding acquisition, Project administration, Supervision, Writing – review & editing.

---

**Fig. S1.**
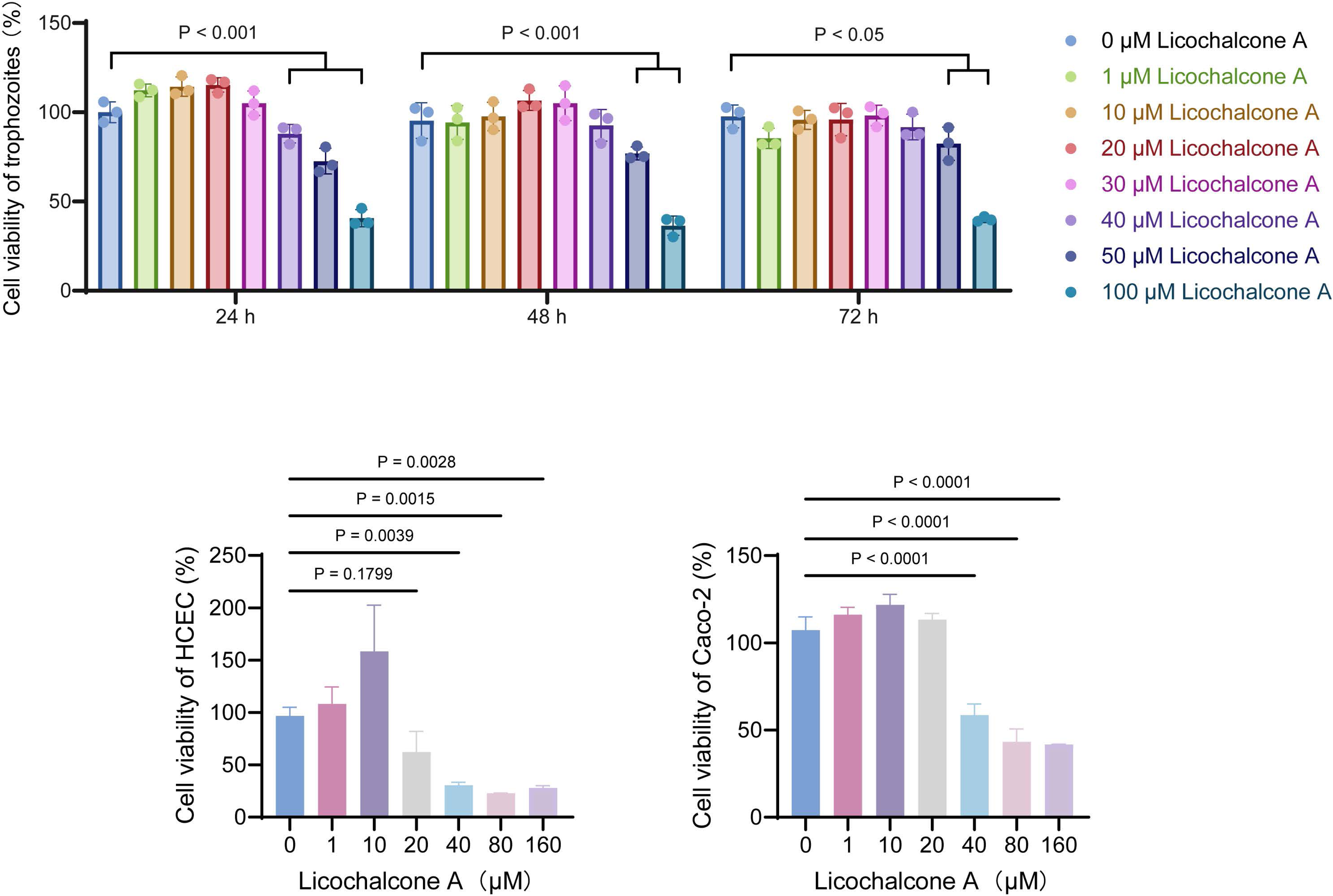
The effect of licochalcone A (Lico A) to *A. castellanii* trophozoites, HCEC and Caco-2 cells.

**Fig. S2.**
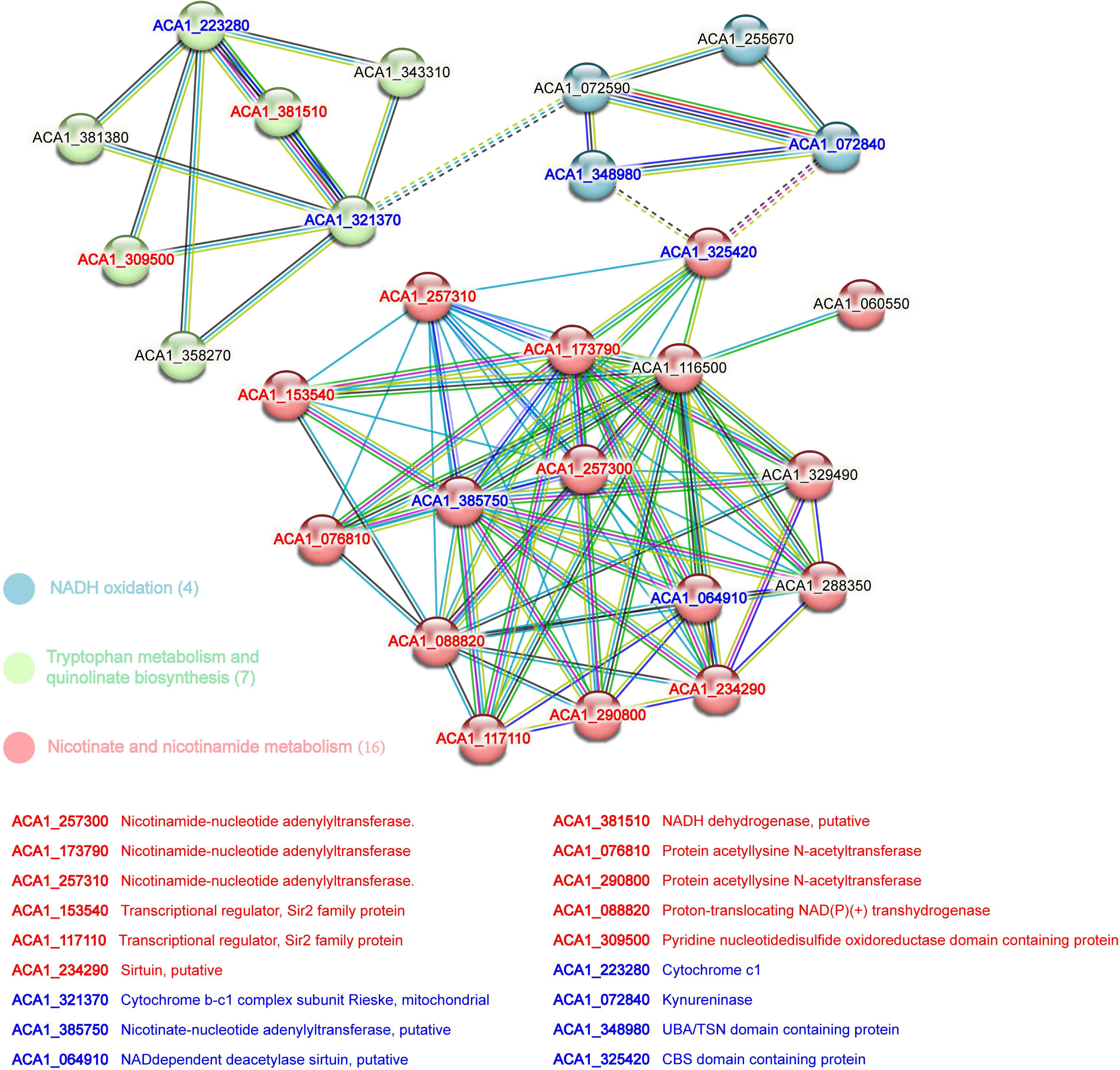
The protein-protein interation (PPI) network constructed using STRING database and RNA sequencing results obtained under 55 μM ISL treatment. Red gene name indicates significant upregulation, blue indicates significant downregulation, while black denotes no change or undetected expression.

**Fig. S3.**
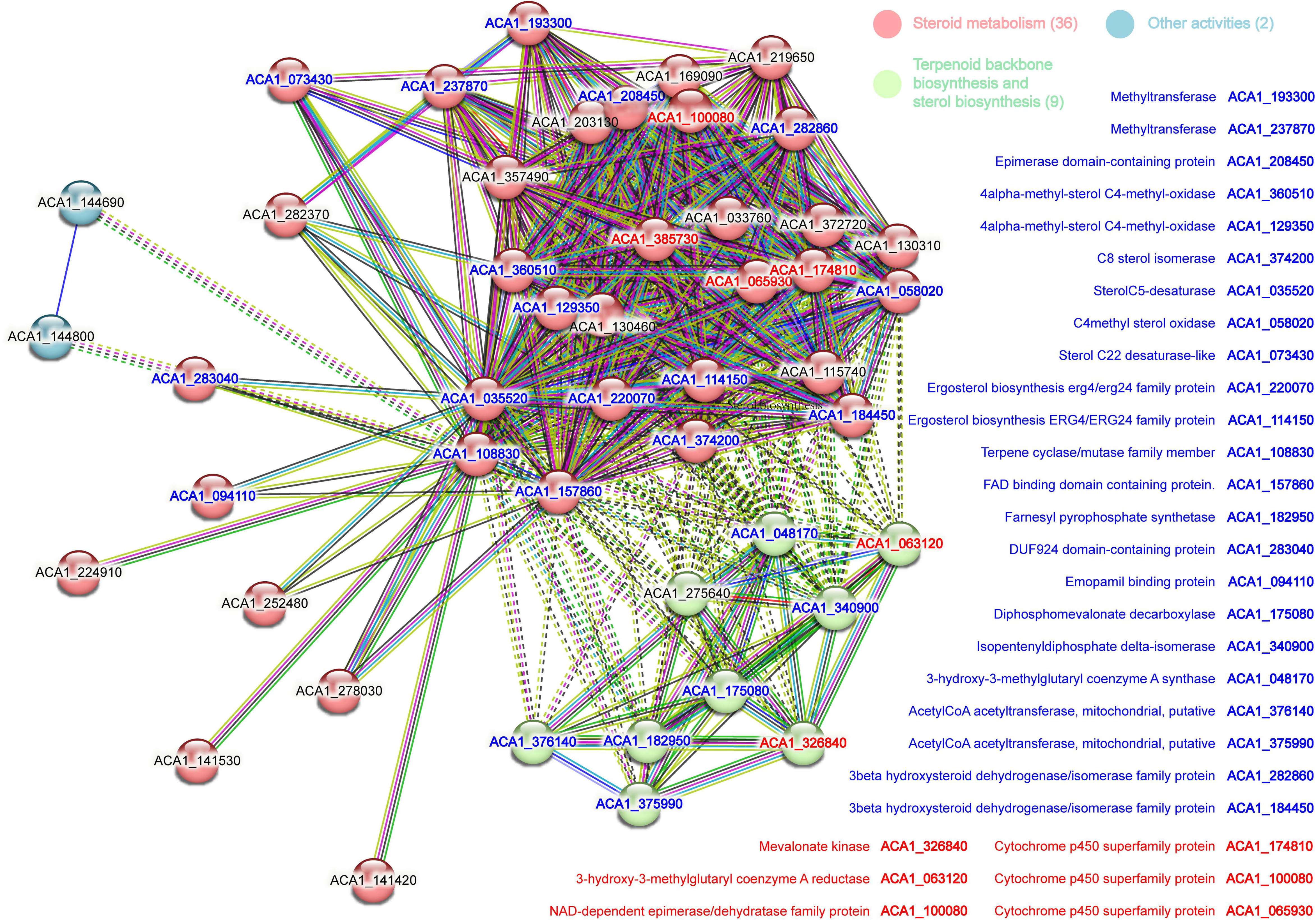
The protein-protein interation (PPI) network constructed using STRING database and RNA sequencing results obtained under 8 μM GLA treatment. Red gene name indicates significant upregulation, blue indicates significant downregulation, while black denotes no change or undetected expression.

