## Supplementary material for "Amoebicidal Action of Licorice-Derived Compounds: Mechanisms and Effects Against *Acanthamoeba castellanii*": Table S1

| Name | Primer | Sequence（5′–3′） | Reference |
| --- | --- | --- | --- |
| 18S | Forward  Reverse | TCCAATTTTCTGCCACCGAA  ATCATTACCCTAGTCCTCGCGC | (Chen et al., 2025) |
| PARP | Forward  Reverse | CGACATCCGCTGGGAGGACAAG  CGCCTGCCTCCACGAAGTTGTC | XM_004333471.1 |
| NAMPT | Forward  Reverse | CTCAAGGTCATCCGCAACGA  GGGGAGATGGGGTCGAAAAC | XM_004333612.1 |
| NMNAT | Forward  Reverse | ATATTTGAGGCGGCGAAGGA  CACGATGAAACCTCGACCCA | XM_004333516.1 |
| GST | Forward  Reverse | ACAAGCCCGATTGGTACTGG  GGAGTAACCTGCGACCCAAA | XM_004343605.1 |
| Sir2 | Forward  Reverse | TACGACCTCCATCCGACCTT  CCGAAGCTAAGGGTGTGCTT | XM_004358188.1 |
| NQO1 | Forward  Reverse | GTGAACCCTTTGGACGCTGA  CGGGTGTGTGGGGTAGATTG | XM_004337797.1 |
| SC5D | Forward  Reverse | CTGCCGTTCCTGATCCTGAC  CGAGGTAGGTCCACTTGTGC | XM_004348686.1 |
| ACAT | Forward  Reverse | ATTTCGCTGGACCGTCTGAA  CAGATCGACGCCACACCAAT | XM_004353631.1  XM_004353628.1 |
| ERG4/ERG24 | Forward  Reverse | CCAACGAGTTTGAGACGCAC  GCATACTGGTGGTGGACGAT | XM_004342119.1 |
| SC4MO | Forward  Reverse | CTCTACAAGCGGTGCCTCAA  GTTCGGATGCCAAGCAGTTC | XM_004339111.1 |
| ERG5-like | Forward  Reverse | ATACCTCGCAGTCGGTGTTC  TCCAGAAGTCCAACAGGCAC | XM_004353175.1 |
| CYP51 | Forward  Reverse | GGCGAGAACTTTGCCTACCT  GTGTACCTGAGAAGGCACGG | XM_004334246.1 |
| GAPDH | Forward  Reverse | TCACCACCATGGAGAAGGC  GCTAAGCAGTTGGTGGTGCA | NM_001256799 |
| APAF-1 | Forward  Reverse | GTCACCATACATGGAATGGCA  CTGATCCAACCGTGTGCAAA | NM_181868 |
| ATM | Forward  Reverse | CAGGGTAGTTTAGTTGAGGTTGACAG  CTATACTGGTGGTCAGTGCCAAAGT | NM_000051 |
| BCL-2 | Forward  Reverse | GGTGGGGTCATGTGTGTGG  CGGTTCAGGTACTCAGTCATCC | NM_000657 |
| BAX | Forward  Reverse | CAGCAAACTGGTGCTCAAGG  CGGAGGAAGTCCAATGTCCA | NM_138763 |
| TNF-α | Forward  Reverse | TAGGCTGTTCCCATGTAGCC  CAGAGGCTCAGCAATGAGTG | NM_000594 |
| TGF-β | Forward  Reverse | CTAATGGTGGAAACCCACAACG  TATCGCCAGGAATTGTTGCTG | NM_000660 |
| FAS | Forward  Reverse | CAGAAGATGTAGATTGTGTGAT  CTTGGTGTTGCTGGTGAGTGTG | NM_152871 |
| SOD2 | Forward  Reverse | GGAAGCCATCAAACGTGACTT  CCCGTTCCTTATTGAAACCAAGC | NM_000636 |

**Table S1. Primer sequences used for real-time PCR.**
